# The Two-Step Cell Death Induction by the New 2-Arachidonoyl Glycerol Analog and its Modulation by Lysophosphatidylinositol in Human Breast Cancer Cells

**DOI:** 10.1101/2024.12.01.626225

**Authors:** Mikhail G. Akimov, Natalia M. Gretskaya, Eugenia I. Gorbacheva, Nisreen Khadour, Galina D. Sherstyanykh, Vladimir V. Bezuglov

## Abstract

2-arachnadoyl glycerol (2-AG) is one of the most common endocannabinoid molecules with antiproliferative, cytotoxic and pro-proliferative effects on different types of tumors. Typically, it induces cell death via CB1/CB2-linked ceramide production. In breast cancer, ceramide is counterbalanced by the sphingosine-1-phosphate, and thus the mechanisms of 2-AG influence on proliferation are poorly understood. In this paper, we evaluated the mechanism of the cell death induction by 2-AG and the influence of LPI on it in 6 human breast cancer cell lines of different tumor degree using a novel analog, 2-arachidonoyl-difluoropropanol (2-ADFP). 2-ADFP induced cell death via the CB2 and TRPV1-dependent COX-2 induction and oxidized metabolites production. The activity was enhanced by LPI.

## 1. Introduction

2-arachnadoyl glycerol (2-AG) is one of the most common endocannabinoid molecules and is usually produced on demand by mesenchymal stromal cells in the bone marrow [1]. Degradation of 2-AG is predominantly mediated by monoacylglycerol lipase (MAGL) or α/β-hydrolase domain containing 6 (ABHD6) and 12 (ABHD12) [2]. Receptors activated by 2-AG are represented by the G_i/o_ protein-bound cannabinoid receptors CB1 and CB2 [3,4]. Affinity for CB2 in 2-AG is greater than for CB1. The vanilloid receptor TRPV1 is an ion channel that can also be activated by 2-AG at concentrations of tens to hundreds of micromoles [5].

In the context of cancer cells, 2-AG has antiproliferative, cytotoxic and pro-proliferative effects on different types of tumors, and is also able to stimulate or, conversely, inhibit cell migration.

Cell migration is inhibited by 2-AG through activation of the CB1 receptor, followed by inhibition of adenylate cyclase activity. This can lead to disruption of the signal transmission pathway to protein kinase A, which regulates cell metastasis. This effect of 2-AG was manifested in micromolar concentrations [6]. Activation of cell migration occurs through the CB2 receptor, resulting in phosphorylation of ERK1/2, which leads to an increase in cell mobility of B-cell leukemia. In this case, micromolar concentrations (microns) are also needed to activate the signaling pathway [7,8].

The antiproliferative effect of 2-AG is typically realized through the CB1 receptor, which is known to inhibit breast cancer cells proliferation [9]. This way, 2-AG reduces viability and induces apoptosis in DU-145 [10], MCF-7 [11], and in part in the Hodgkin’s lymphoma cells [12].

However, the cytotoxic effect on the tumor cells of the larynx HNSCC and oral cavity SNU-1066 is associated with the activation of the CB2 receptor [13]. In addition, CB2 agonist induced cytotoxicity and apoptosis in human colorectal cancer cells (HT-29) [14], and the selective CB2 agonist JWH-133 induced a considerable regression of malignant tumors generated by inoculation of C6 glioma cells [15].

The mechanisms by which the CB1 receptor activation reduces proliferation and induces apoptosis are rather well-known and typically occurs via the ceramide biosynthesis stimulation [16]. CB2 activation may also lead to ceramide biosynthesis as a part of the apoptosis induction [17]. Ceramide, in part, is able to induce endoplasmic reticulum stress and subsequent apoptosis [18]. However, in the breast cancer setting, ceramide biosynthesis induction may cause a pro-survival effect due to high rate of conversion to sphingosine-1-phosphate [19]. In addition, CB2 activation in breast cancer cells was shown to modulate viability in a calcium-dependent fashion [20]. In addition, a prolonged treatment by the CB2 agonists could have a pro-survival effect on cancer cells [21]. These data suggest the existence of multiple mechanisms, by which 2-AG could induce cell death, especially in the breast cancer setting, and their elucidation could be important for the understanding of the role of this substance in cancer development.

In addition to autonomous effects, the 2-AG CB2 receptor is involved in interreceptor interaction. As such, heterodimers of CB2 with the chemokine receptor CXCR4 [22], adenosine receptor (A2aR) [23], and HER2 [24] were described. Of special interest are the heterodimers with other cannabinoid receptors, as they could lead to an unexpected interaction between various endocannabinoids. As such, in the CB1-CB2 complex, CB2 blocks the effects mediated by the CB1 receptor [25]. GPR55-CB2 hetero- dimers were also identified [26], pointing to the possibility of interaction between the pro-cancer LPI signaling axis [27] with the 2-AG function.

In this paper, we evaluated the mechanism of the cell death indication by 2-AG in 6 human breast cancer cell lines of different tumor degree, and the influence of LPI on it. 2-AG induced cell death via the CB2 and TRPV1-dependent COX-2 induction and oxidized metabolites production. The activity was enhanced by LPI.

### 2. Results

### 2.1. Cell Lines

The topic of this research was the direction and mechanism of 2-AG and LPI combined activity. As with the case of anandamide in our previous work [28], we assumed that at least the direction and magnitude of this interaction could depend on the grade of a tumor, as the cells progressively accumulate mutations and their response to various stimuli change. To address this possibility, a panel of human breast cancer cell lines with different properties was used (Table 1). These lines were chosen to represent all of the major breast cancer subtypes [29,30].

**Table 1.** Subtypes and major receptor expression patterns in the cell lines used [29,30].

| Cell Line |  |  |  |  |  |  |
| --- | --- | --- | --- | --- | --- | --- |
|  | MCF-10A | MCF-7 | BT-474 | SK-BR-3 | BT-20 | MDA-MB-231 |
| Grade | Non-transformed | Luminal A | Luminal B | HER2 | Basal | Basal |
| Metastasis | - | Non-invasive [31] | Low [32] | Very low [32] | Low | ++ |
| ER | - | + | + | - | - | - |
| PR | - | + | + | - | - | - |
| HER2 | - | - | + | + | - | - |

### 2.2. Synthesis and Characterization of the 2-AG Analog 2-Arachidonoyl-1,3-difluoropropanol

Given the low stability and complicated chemical synthesis of 2-AG, its analog 2-arachidonoyl-1,3-difluoropropanol (2-ADFP) with hydroxyl groups replaced by the fluorine was used (Figure 1).

**Figure 1.**
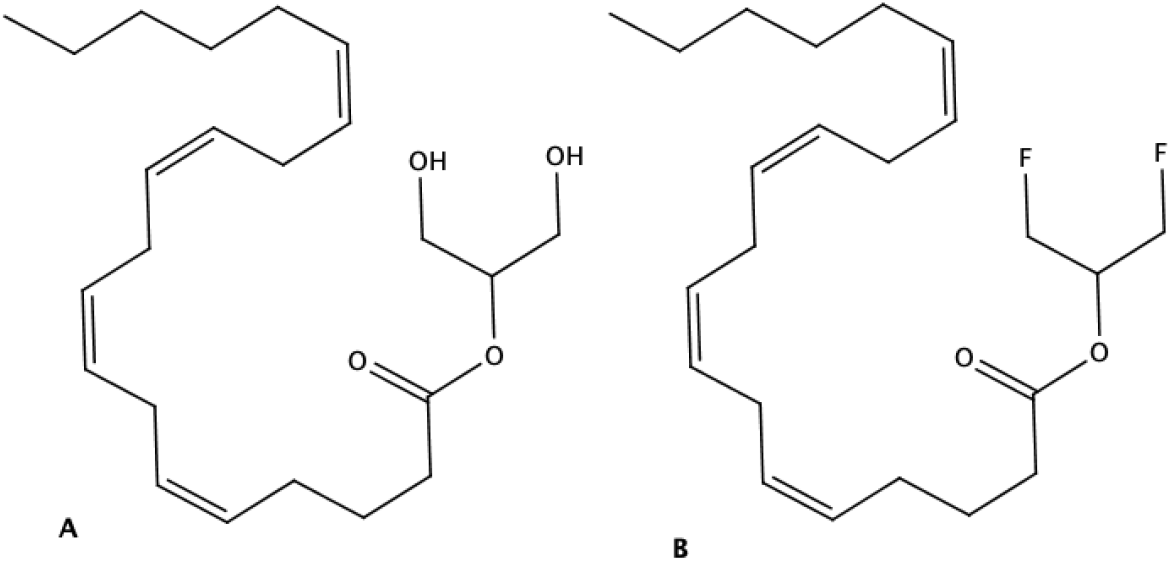
The structure of 2-AG (**A**) and 2-ADFP (**B**)

To validate the molecular target of 2-ADFP, we used a molecular docking approach and a cellular model. First, the affinity of the molecule to the active site of the CB2 and CB1 receptor were estimated in the molecular docking experiments in comparison to the unmodified 2-AG (Table 2). The affinity scores of the substances were very close to each other, however, 2-ADFP demonstrated a stronger affinity towards CB1 in the antagonist-bound conformation.

**Table 2.** AutoDock Vina affinity scores of 2-ADFP and 2-AG towards the CB1 and CB2 receptor in the molecular docking experiments. CB1: an antagonist-bound receptor PDB ID 5TGZ [33] and an inverse agonist-bound receptor 5U09 [34], CB2: an agonist-bound receptor PDB ID 6KPC [35].

|  | Receptor |  |  |
| --- | --- | --- | --- |
|  | CB1<br>Inverse<br>Agonist-bound | CB1<br>Antagonist-bound | CB2<br>Agonist-bound |
| Affinity score |  |  |  |
| 2-ADFP | -7.757 | -8.076 | -8.687 |
| 2-AG | -7.685 | -7.564 | -8.638 |

Next, we tested 2-ADFP activity on the DU 145 cell line, for which a substantial expression of the CB2 receptor, the primary target of 2-AG, is known [36]. As a control, we silenced the receptor using siRNA (Figure 2).

**Figure 2.**
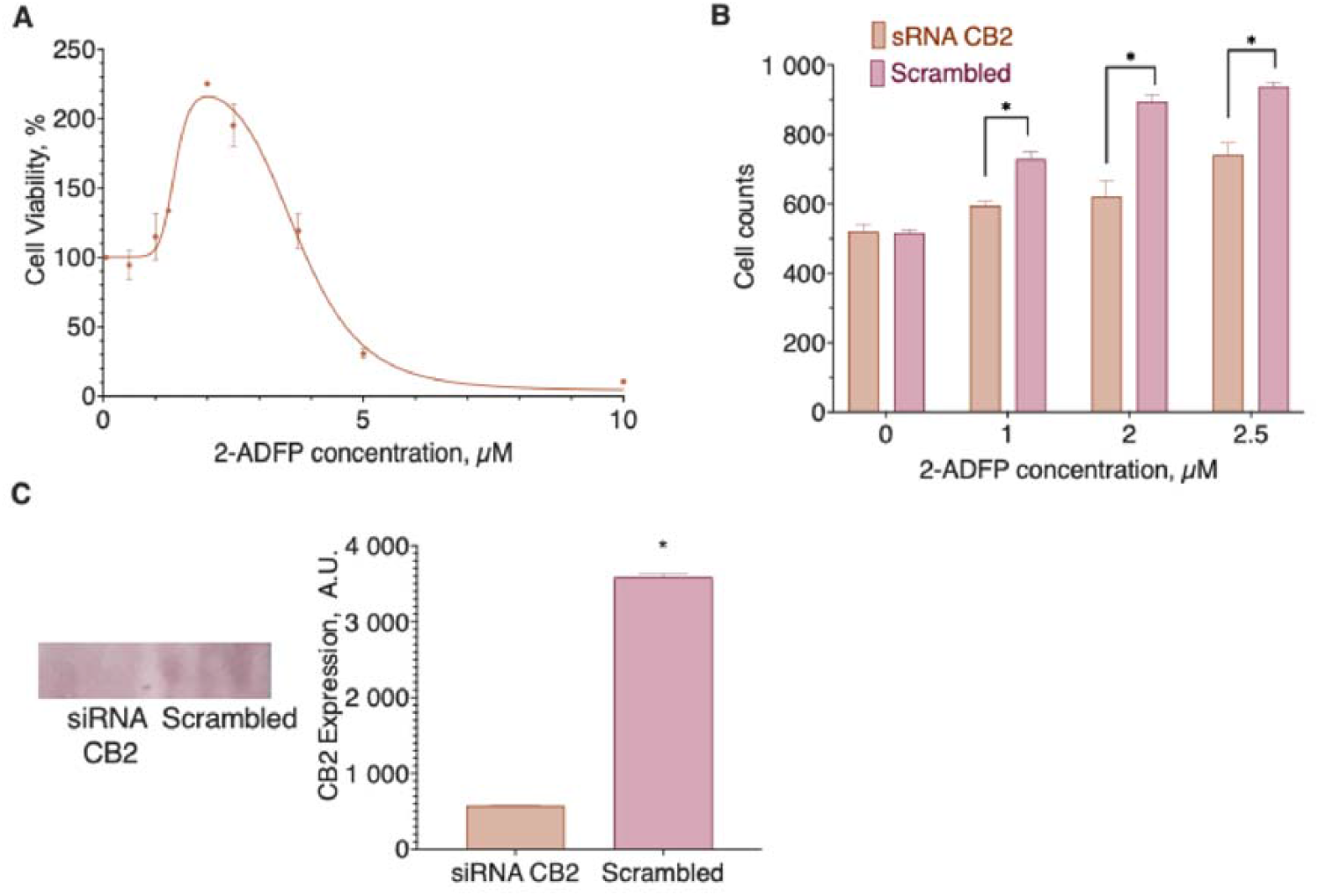
CB2 receptor role in the DU145 cell line response to the 2-ADFP action. (**A**) The influence of 2-ADPG concentration on the cell viability, (**B**) the effect of the CB2 receptor knockdown on the pro-proliferative 2-ADFP activity, (**C**) siRNA knockdown of the CB2 receptor in the DU 145 cell line. Incubation time 72 hours, resazurin test, mean±standard error (N=4 experiments). *, a statistically significant difference from the cells transfected with the scrambled siRNA, p≤0.05, ANOVA with Holm-Sidak post-test (**B**) and Student’s t-test **(C**)

2-ADFP stimulated the cell proliferation in the concentration range from 1 to 3 µM (EC_50_ 1.37 (1.023 to 1.717) µM), and this activity was abolished after the CB2 receptor knockdown, thus confirming the ability of the substance to interact with the receptor. The higher concentrations of the substance were cytotoxic.

### 2.3. Individual Effects of 2-ADFP on the Breast Cancer Cell Lines

The first stage of the research was to determine the individual activity of 2-ADFP for the chosen cell lines. As far as 2-AG is known to affect cell proliferation [37,38], and LPI can act as a proliferation stimulator, the incubation time was chosen to be 72 h. Cell viability was determined using the resazurin test; cell death was confirmed using microscopy.

We did not explicitly test LPI activity, as it was already described in our previous work [28].

In the range from 1 to 50 µM, 2-ADFP did not affect the proliferation of the breast cancer cell lines. However, in the concentration range from 50 to 150 µM, the substance was cytotoxic with EC_50_ in the range 89-167 µM (Figure 3, Table 3). For the MCF-10A, MCF-7, and BT-474, the highest concentration of the substance (200 µM) did not kill 100% of the cells, while for the SK-BR-3, BT-20, and MDA-MB-231, all of the cells died at the high substance concentrations.

**Table 3.** The effect of 2-ADFP on breast cancer cell lines proliferation. Incubation time 72 hours, resazurin test, mean±standard error (N=4 experiments).

|  | Cell Line |  |  |  |  |  |
| --- | --- | --- | --- | --- | --- | --- |
|  | MCF-10A | MCF-7 | BT-474 | SK-BR-3 | BT-20 | MDA-MB-231 |
|  | EC<sub>50</sub>, μM, mean (95% CI) |  |  |  |  |  |
| 2-ADFP | 143.6<br>(140.6-146.6) | 149.7<br>(143.3-156.3) | 167.3<br>(156.8-177.7) | 89.98<br>(83.34-97.1) | 97.80<br>(110.7-112.3) | 118.2<br>(116.0-120.7) |

**Figure 3.**
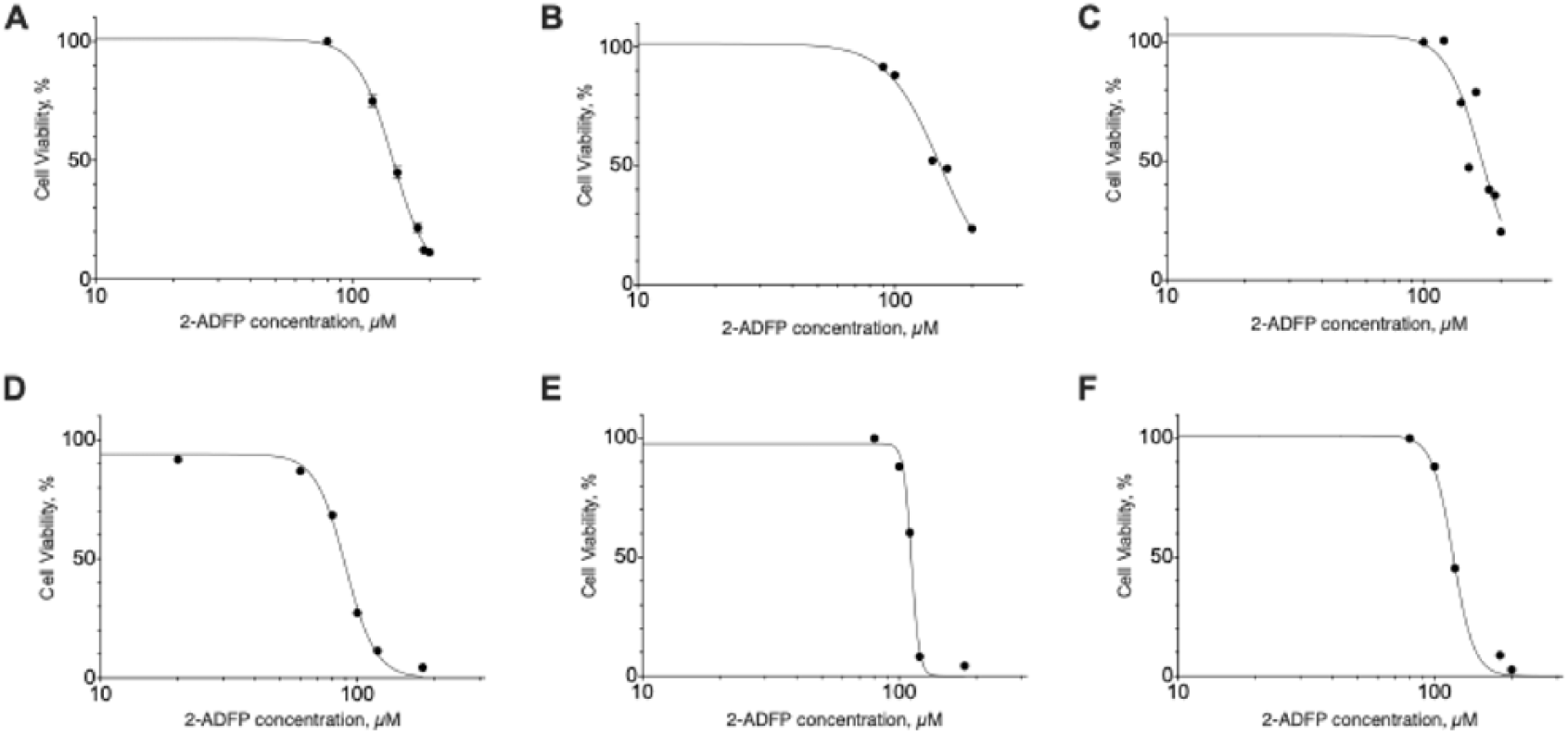
The effect of 2-ADFP on breast cancer cell lines proliferation. Incubation time 72 hours, resazurin test, mean±standard error (N=4 experiments). (**A**) MCF-10A, **(B**) MCF-7, (**C**) BT-474, (**D**) SK-BR-3, (**E**) BT-20, (**F**) MDA-MB-231

### 2.4. The Effect of 2-ADFP-LPI Combinations on the Viability of the Cell Lines

We next tested the ability of LPI to change the activity of 2-ADFP when added together. We used a 2-ADFP concentration close to its EC_50_ value and several LPI concentrations to account for the possible activity changes in different concentration ranges. In this experiment series, we also used a 72-h incubation with the substances to detect possible proliferation changes and resazurin test to evaluate cell viability.

In the case of the MCF-10A cell line, LPI did not affect the 2-ADFP activity, while in the case of all other cell lines the combined cytotoxicity was higher than the one of the 2-ADFP alone (Figure 4).

**Figure 4.**
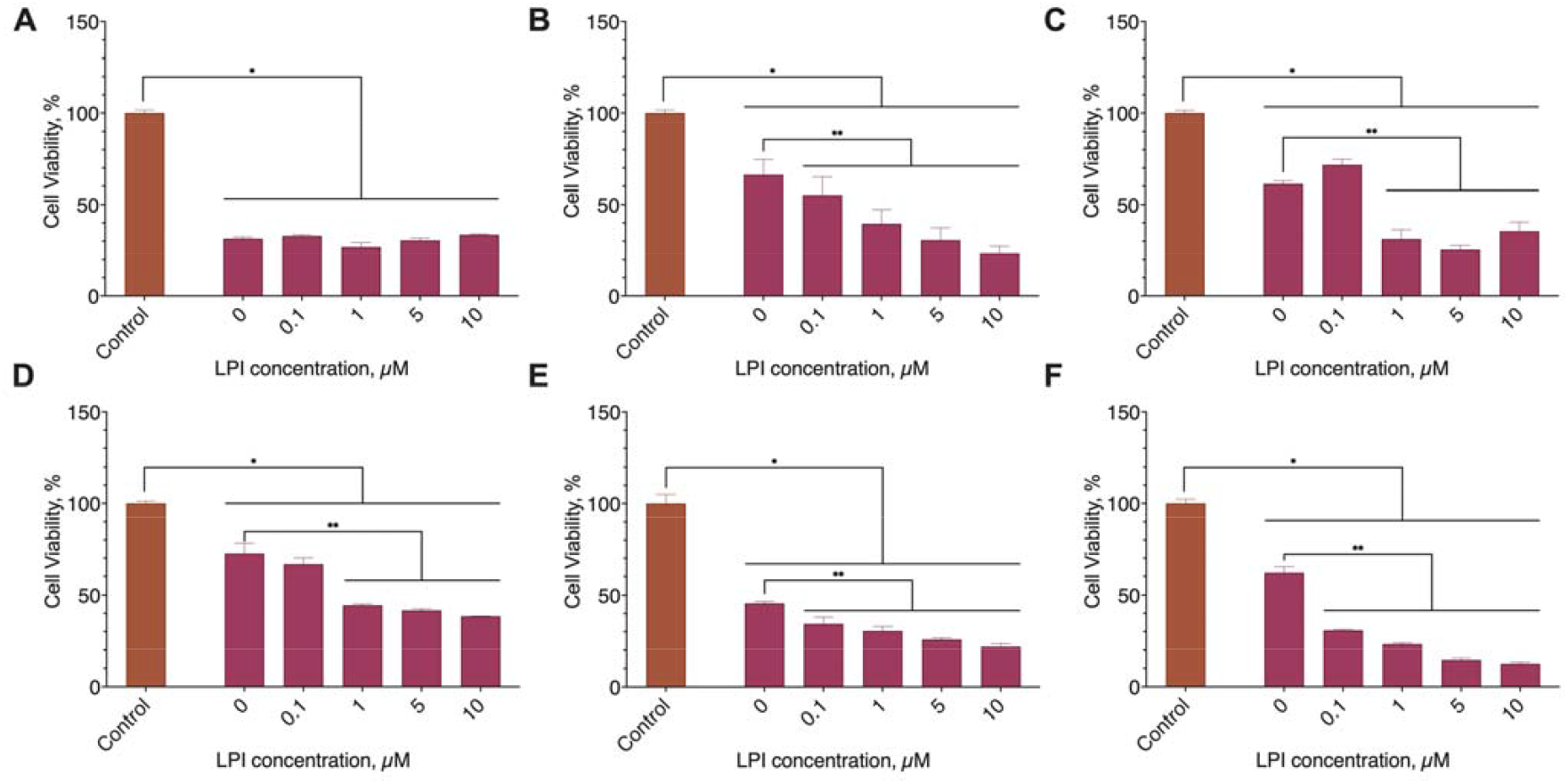
The effect of 2-ADFP combination with LPI on breast cancer cell lines viability. Incubation time 72 hours, resazurin test, mean±standard error (N=4 experiments). (**A**) MCF-10A, (**B**) MCF-7, (**C**) BT-474, (**D**) SK-BR-3, (**E**) BT-20, (**F**) MDA-MB-231. **, a statistically significant difference from 2-ADFP alone, p≤0.05, ANOVA with Holm-Sidak post-test; *, a statistically significant difference from the non-treated control, p≤0.05, ANOVA with Holm-Sidak post-test

### 2.5. Receptor Participation in the Individual and Combined Substance Effects

The next question of our research was the mechanism of the LPI effect on the 2-ADFP activity. Our hypothesis was that 2-ADFP-LPI interaction could proceed through one of the known cannabinoid receptors, as some of them (CB1 and CB2) are targets for 2-ADFP [39], others are targets for LPI (GPR18 and GPR55) [40], and at least for some of them, a formation of activity-changing heterodimers was described [41–43].

The expression of the core cannabinoid receptors (CB1, CB2, and GPR55) in the model cell lines was estimated in our previous work. MCF-7, MDA-MB-231, SK-BR-3, BT-474, and BT-20 expressed all three of them, although to a different extent. In MCF-10A, the expression of CB2 was not observed, and in MCF-7, GPR55 expression was negligible [28].

We added a selective blocker for each of those receptors both to 2-ADFP alone, and to the 2-ADFP-LPI combination to check for the importance of the appropriate receptor, and evaluated the proliferation change in the resazurin test after 72 h of incubation with the cells.

By their response to the 2-ADFP treatment the cell lines could be separated into three groups (Figure 5):

**Figure 5.**
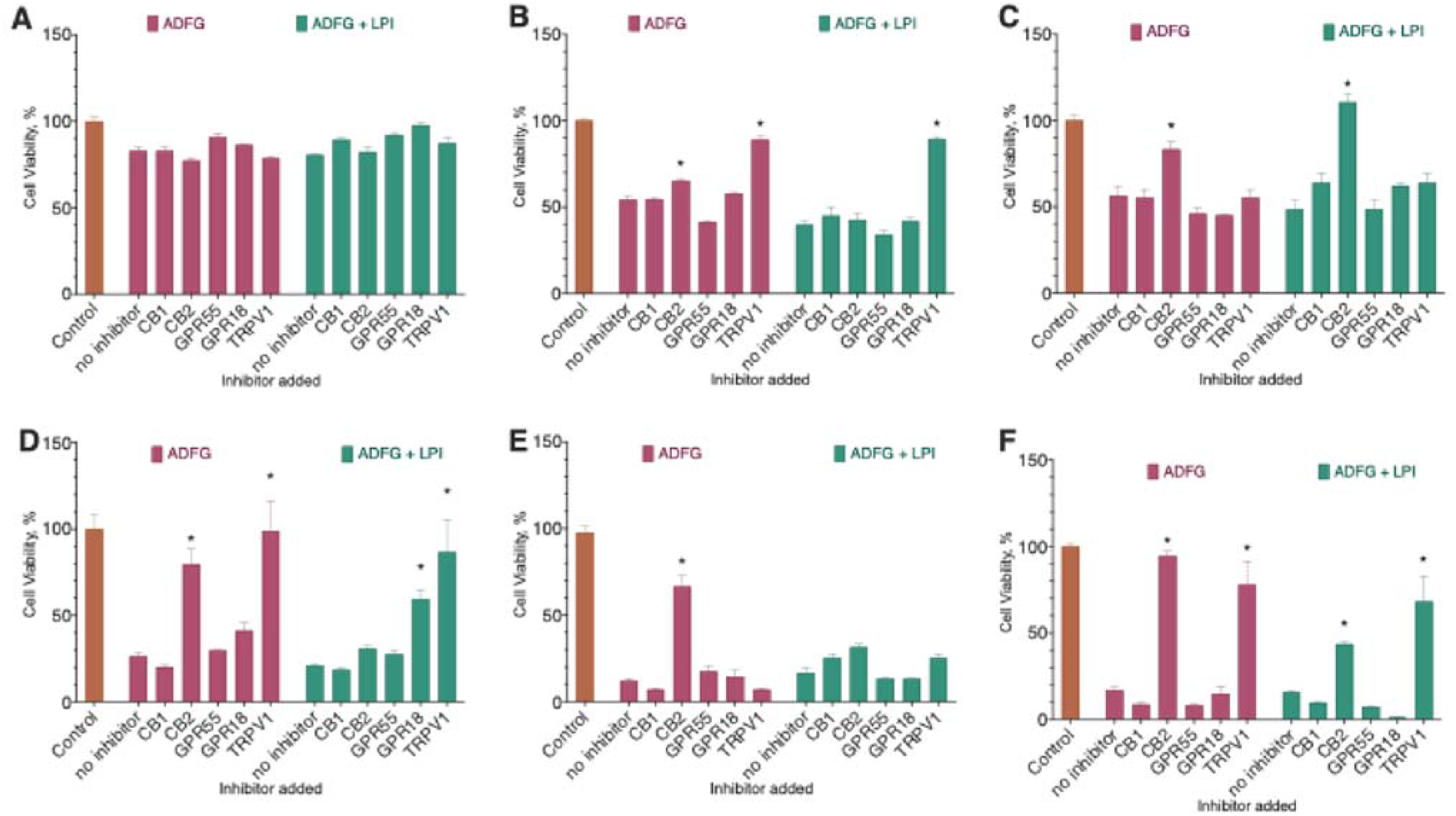
The effect of receptor blockers on the effect of 2-ADFP (EC_50_-EC_70_) combination with LPI (5 µM) on breast cancer cell lines viability. The following substances and concentrations were used: CB1, SR 141716A (100 nM); CB2, SR 144528 (100 nM); GPR55, ML-193 (2 µM); GPR18, PSB CB5 (3 µM); TRPV1, capsazepine (5 µM). Incubation time 72 hours, resazurin test, mean±standard error (N=4 experiments). (**A**) MCF-10A, (**B**) MCF-7, (**C**) BT-474, (**D**) SK-BR-3, (**E**) BT-20, (**F**) MDA-MB-231. *, a statistically significant difference from 2-ADFP alone, p≤0.05, ANOVA with Holm-Sidak post-test

- MCF-10A: neither of the receptor blockers used caused a substantial reduction of the substance activity.
- BT-474 and BT-20: the CB2 receptor blocker substantially decreased 2-AG activity.
- MCF-7, SK-BR-3, and MDA-MB-231: both CB2 and TRPV1 receptor blockers substantially decreased the 2-AG activity.

In most cases, the addition of LPI shifted the response from CB2 towards TRPV1. These data point to the participation of these receptors in the response to 2-ADFP.

### 2.6. Signal Transduction Downstream the CB2 Receptor

At the first glance, the observed participation of both of TRPV1 and CB2 receptors in the cytotoxic activity of 2-ADFP seemed somewhat counterintuitive. However, the unifying activity of these receptors is the increase of the intracellular Ca^2+^ [44,45]. The increase of Ca^2+^ could lead to the activation of the CaMKII/CREB pathway, which in turn stimulates the expression of the COX-2 enzyme [46]. Once produced, it can oxidize the arachidonic moiety of 2-AG to PGE_2_, which is able to induce apoptosis [47]. To validate the participation of this pathway, we used several inhibitors to block some of the key signal transduction events: diclofenac (COX-2, 3 µM), N-acetylcysteine (NAC, removes oxidative stress linked to the PGE_2_-glycerol toxicity [47], 10 µM), Xestospongin C (IP3R, 3 µM), and 666-15 (CREB, 3 µM). As the model, we used MDA-MB-231 cell line with a known endogenous COX-2 expression, and BT-20, which does not express COX-2 [48].

In both cell lines, NAC and diclofenac substantially reduced the cytotoxic effect of 2-ADFP, indicating the crucial role of the enzyme (Figure 6). Xestospongin C was also active in both cell lines, however, in MDA-MB-231 cell line it only reduced the cytotoxicity by about 50%. This agrees with the data on the participation of TRPV1 receptor in the 2-ADFP signaling in this cell line.

**Figure 6.**
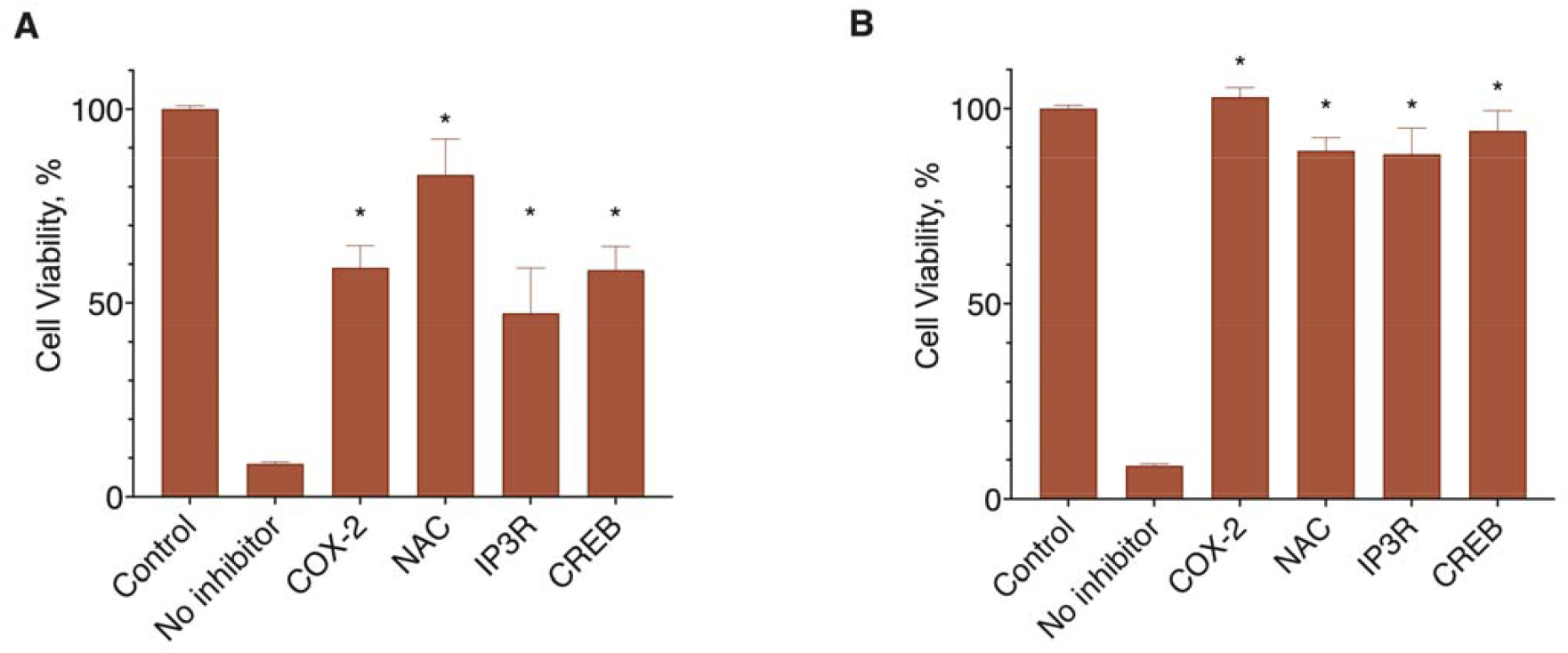
The participation of the COX-2 in the 2-ADFP cytotoxicity. (**A**) MDA-MB-231, (**B**) BT-20. Incubation time 72 hours, resazurin test, mean±standard error (N=4 experiments). *, a statistically significant difference from 2-ADFP without inhibitor, p≤0.05, ANOVA with Holm-Sidak post-test

## 3. Discussion

In this paper, we studied the effect of 2-ADFP and its modulation by LPI in a panel of human breast cancer cell lines of different tumor grade. This interaction could be quite important in the cancer setting, as the endocannabinoid system is a possible target for anti-cancer therapy [49], and LPI is usually considered a pro-proliferative molecule [50]. We found that depending on the receptor set of the cell line, 2-ADFP effect could occur via different receptor targets, and usually it is enhanced by the LPI presence.

At the first stage of the work, we developed an easy to synthesize analog of 2-AG by replacing its hydroxyl groups by fluorine. This molecule could be easily obtained by the F-anhydride method [51]. Molecular docking experiments together with the data of the CB2 receptor knockdown experiments demonstrated, that 2-ADFP should have similar affinity for the CB2 receptor, but in the case of the CB1 receptor it behaves more like an antagonist. Thus, we did not expect to observe any CB1-based effects for this compound.

2-ADFP induced cell death in all six cell lines studied. The obtained EC_50_ values (89-167 µM) were relatively low compared to the typical data on the 2-AG activity from the literature [52], but were in the line with our data on the low CB2 receptor expression on these cell lines [28]. On the other hand, for the DU 145 cells, which express larger quantities of CB2, the EC_50_ (1.37 µM) value was close to the ones typically observed in the literature. The addition of LPI resulted in the dose-dependent enhancement of the 2-ADFP cytotoxicity, but only for the malignant cell lines. This behavior was rather unexpected, as the typical activity of LPI is the stimulation of the proliferation [27]. On the other hand, we have already observed a similar interaction pattern for the LPI-anandamide pair [28].

Our next aim was to reveal the mechanisms of the observed cytotoxic effects. As far as both 2-AG and LPI belong to the endocannabinoid system, we tried to block their activity using the selective blockers for the classical cannabinoid receptors CB1 and CB2, vanilloid receptor TRPV1, and non-classical cannabinoid receptors GPR55 and GPR18. In these experiments, only the blockers for the CB2 and TRPV1 receptors were active, and the addition of LPI shifted the effect towards less sensitivity to the CB2 receptor inhibition. The participation of CB2 and TRPV1 receptors in the 2-ADFP response seems to be logical, as both of these receptors recognize 2-AG [3,4,53]. The absence of the CB1 response contradicts the literature data on the mechanisms of the anti-proliferative effect of 2-AG [10–12]. However, given our data on the affinity scores of 2-ADFP towards different CB1 receptor conformations, it could probably be due to the antagonism of 2-ADFP for the CB1 receptor. The lack of the response to the GPR55 and GPR18 receptor blockers in the experiments on the LPI activity could be due to the participation in this interaction of its other receptor GPR119 [54].

In the context of the breast cancer cell lines, a CB2-based cytotoxicity is a rather unexpected behavior, as the main signal route for it, ceramide biosynthesis, is counterbalanced by the sphingosine-1-phosphate synthesis in this setting [19]. However, 2-AG could induce apoptosis indirectly, after the oxidation to its PGE_2_ derivative [47]. We checked this possibility by the addition of the inhibitor of the involved oxidase, diclofenac. Diclofenac fully protected cells against the 2-ADFP cytotoxicity, indicating the involvement of this pathway. However, based on the literature data [48], basal expression of COX-2 is only observed for the MDA-MB-231 cell line. On the other hand, both CB2 and TPRV1 can lead to the increase of Ca^2+^ concentration in the cells, and this event via the CaMKII/CREB activation could induce the COX-2 expression. To check for the participation of this pathway, we used the IP3R blocker (CB2 activates PLC, which produces IP3, and this ligand in turn activates and opens IP3R Ca^2+^ channel [46]), and CREB inhibitor 666-15. Both substances prevented the cytotoxicity of 2-ADFP, confirming the hypothesis. The calcium-dependent activation of COX-2 expression is in the line of the possible activation of the GPR119 receptor by LPI, as this protein could also induce Ca^2+^ mobilization [55].

The resultant sequence of events during the cell death induction by 2-ADFP and its modulation by LPI is summarized on the Figure 7.

**Figure 7.**
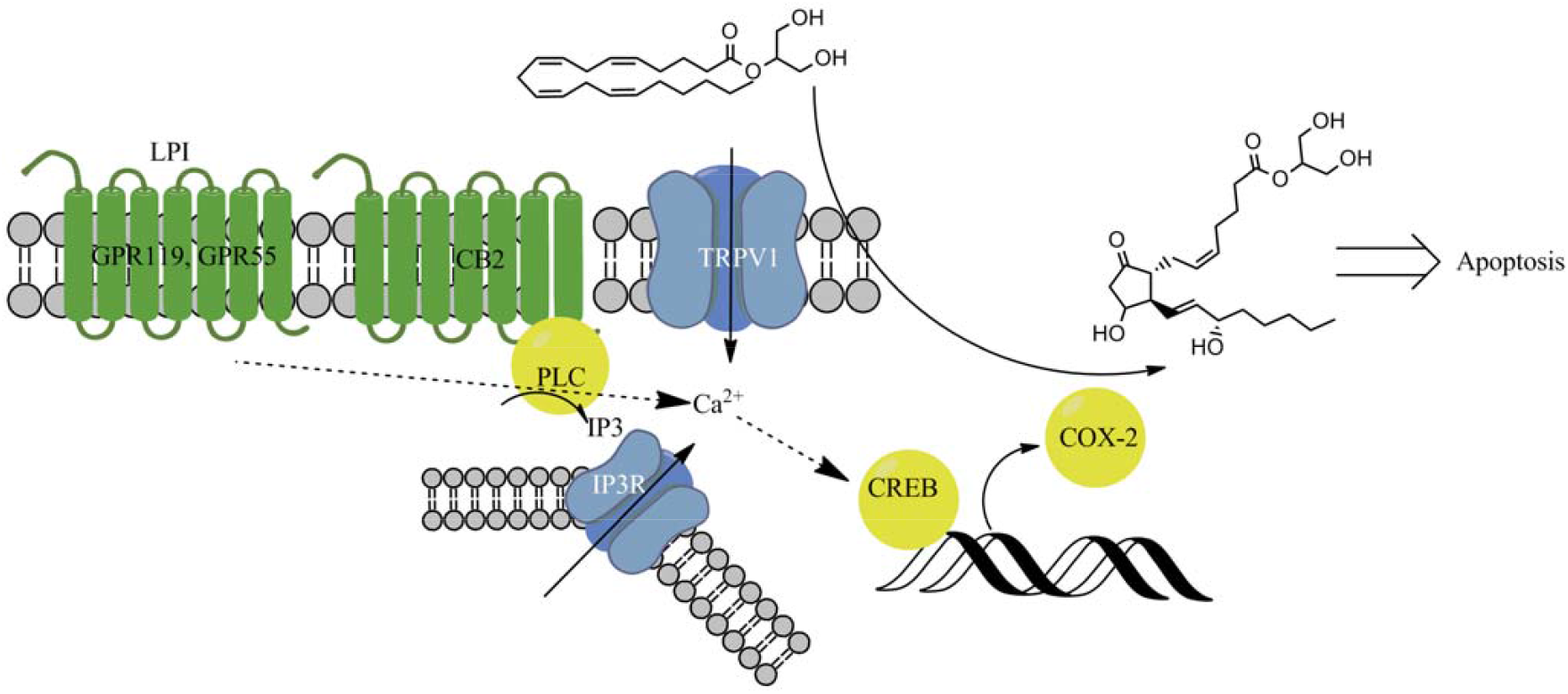
Signaling during the CB2/COX-2-dependent cell death induction by 2-AG.

The observe two-stage mechanism of 2-AG cytotoxicity is an interesting addition to the already known mechanisms of the activity of this compound. On one hand, it could be used as a novel principle for the rational design of anti-cancer compounds. On the other hand, the oxidation products produced by COX-2 may have pro-proliferative activity [56], and so the discovered COX-2 induction and 2-AG metabolic transformations should be taken into the account during endocannabinoid-based cancer treatment development.

## 4. Materials and Methods

### 4.1. Reagents and Cell Lines

DMEM/F12, DMEM, L-glutamine, HCl, Earle’s salts solution, Hank’s salts solution, Versene’s solution, antibiotic/antimycotic mixture (penicillin, streptomycin, amphotericin B), DMEM, trypsin, and fetal bovine serum were from Servicebio, Hubei, China.

Cell lines MDA-MB-231 (HTB-26), MCF-10A (CRL-10317), MCF-7 (HTB-22), BT-474 (HTB-20), BT-20 (HTB-19), SK-BR-3 (HTB-30), and DU 145 (HTB-81) were purchased from ATCC, Manassas, VA, USA.

Antibodies anti-b-actin and anti-CB2 were from Abcam, Cambridge, UK. Anti-mouse IgG antibody was from Jackson ImmunoResearch, Cambridge, UK.

SR 144028, PSB CB5, ML-184, ML-193, capsazepine, and LPI was from Tocris Bioscience, Bristol, UK. DMSO, resazurin, D-glucose, glycylglycine, acetic acid, MgSO_4,_ EGTA, dithiothreitol, acrylamide, bis-acrylamide, Triton X-100, SDS, nitro blue tetrazolium, Tris, EDTA, agarose, bicinchoninic acid, bovine serum albumin, anti-rabbit IgG antibody, diclofenac, N-acetyl cysteine, and 5-Bromo-4-chloro-3-indolyl phosphate-toluoidine were from Sigma-Aldrich, St. Louis, MO, USA. The purity of all used reagents was 95% or more.

### 4.2. Chemical Synthesis

Conjugate of arachidonic acid and 1,3-difluoro-2-propanol (2-ADFP) was prepared by the F-anhydride method [51]. Briefly, 150 mg (0.5 mmol) of arachidonic acid were dissolved in 3 ml of MeCN and 70 µl of Py (0.83 mmol) and 70 µl (0.83 mmol) of cyanuric fluoride were added. The mixture was stirred at 20 C under argon for 60 min. Arachidonic acid fluoride was added to 200 µl (2 mmol) of 1,3-difluoro-2-propanol in 2 ml THF and 60 mg (0.5 mmol) of DMAP was added. The reaction mixture was extracted with EtOAc. The final product was purified on a Kieselgel 60 column, eluting with benzene (117.2 mg, 62% of theoretical). PMR spectrum (δ, ppm, J/Hz): 0,98 (3H, t, H-20), 1,40 (6H, m, H-17,18,19), 1,83 (2H, m, H3), 2,17-2,21 (4H, m, H4,16), 2,5 (2H, m, H2), 2,91 (6H, m, H-7,10, 13), 4,56(2H, d, J=4,57, -CH2- difluoropropanole), 4,79(2H, d, J=4,8 -CH2- difluoropropanole), 5,47 (8H, m, H-5,6,8,9,11,12).

### 4.3. Cell culture

Cells were cultured in DMEM (cell lines MDA-MB-231, BT-474, MCF-7, and BT-20), McCoy 5A (cell line SK-BR-3) or RPMI 1640 (cell line DU 145) medium supplemented with 10% FBS, 4 mM (2 mM in the case of the DU 145 cell ine) L-glutamine, 100 U/ml penicillin, 100 µg/ml streptomycin, and 2.5 µg/ml amphotericin B. The medium for the MCF-10A consisted of DMEM/F12 supplemented with 10% FBS, 4 mM L-glutamine, 100 U/ml penicillin, 100 µg/ml streptomycin, 2.5 µg/ml amphotericin B, 20 ng/ml EGF, 0.5 µg/ml hydrocortisone, 10 µg/ml insulin, and 100 nM (−)-isoproterenol [57]. All the cells were maintained in 5% CO_2_ and 100% humidity at 37°C. Cells were subcultured by successive treatment with Versene’s solution and trypsin solution. Mycoplasma contamination was controlled using the Jena Biosciences kit according to the manufacturer’s instructions.

### 4.8. Cytotoxicity and proliferation evaluation

Cells were plated in 96-well plates in the amount of 2500 per well in a volume of 100 µl of medium and cultured for a day. After that, the substance was added at the required concentration in the range from 0.01 to 200 µM in the form of a DMSO solution in a fresh 100 µl of the medium, in which the serum was replaced with a delipidated analog; the final DMSO concentration was 0.5% or less. If receptor blockers were used, they were added to 50 µl of the medium one hour before the addition of the substance and then the second portion (to ensure the constancy of concentration) with the substance; the volume of the medium in which the substance was added was 50 µl in this case. For the proliferation induction studies, the medium with the delipidated FBS was used to eliminate the influence of the natural LPI. The cells were incubated with the substance for 72 hours, after which the viability was determined using the resazurin test, and the cell death using the LDH test. Each experiment was repeated at least five times.

### 4.9. RNA isolation and cDNA synthesis

RNA was isolated from in vitro cultivated cells using the QIAzol Lysis Reagent (Qiagen, Hilden, Germany) as described in manufacturer’s protocol. RNA samples were treated with DNase I (Thermo Fisher Scientific, Lithuania) according to the manufacturer’s instruction. The RNA concentration was measured using a Nanodrop OneC Spectrophotometer (Thermo Fisher Scientific, USA). cDNA was synthesized with MMLV RT kit (Evrogen, Moscow, Russia) according the manufactures’s recommendations.

### 4.10. qPCR

qPCR was performed on LightCycler 96 (Roche) with qPCRmix-HS SYBR reagent (Evrogen, Moscow, Russia). Cycling conditions were 95°C for 150 s, and then 45 cycles of 95°C for 20 s, 57 °C for 20 s and 72°C 20 s. Primer specificity was confirmed by visualizing DNA on an agarose gel following PCR. The relative level of expression was determined by the 2-ΔΔCt method. XXX was used as an internal control. Each analysis was performed in triplicate. Primer sequences for PCR and qPCR were as follows: beta-2 microglobulin forward 5’-CAGCAAGGACTGGTCTTTCTAT-3’, reverse 5’-ACATGTCTCGATCCCACTTAAC-3’; COX-2 forward 5’-GTGCCTGGTCTGATGATGTATG-3’, reverse 5’- CCTGCTTGTCTGGAACAACT-3’; POL2R forward 5’-CCCAGCTCCGTTGTACATAAA-3’, reverse 5’-TCTAACAGCACAAGTGGAGAAC-3’.

### 4.11. Resazurin Test

To evaluate cells viability, the culture medium in the wells was replaced with a 0.2 mM resazurin solution in Earle’s solution with the addition of 1 g/l D-glucose and incubated for 1.5 hours at 37°C under cell culture conditions [58]. After that, the fluorescence of the solution was determined at the excitation wavelength of 550 nm and the emission wavelength of 590 nm using the Hidex Sense Beta Plus microplate reader (Hidex, Finland). The positive control was the cell culture treated with the solvent alone, and the negative control was treated with 0.9% Triton X-100.

### 4.12. CB2 Receptor siRNA Knockdown

To test the hypothesis about the involvement of CB2 receptor as the 2-ADFP target, the knockdown approach of this receptor using siRNA was used. Three duplex siRNAs were used: 1) sense 5′-CCAGGUCAAGAAGGCCUUUdTdT-3′, ant-sense 5’-AAAGGCCUUCUUGACCUGGdTdT-3’; 2) sense 5′- GCUUGGAUUCCAACCCUAUdTdT-3′, anti-sense 5’-AUAGGGUUGGAAUCCAAGCdTdT-3’; 3) sense 5′-CCUGGCCAGUGUGGUCUUUdTdT-3′, anti-sense 5’-AAAGACCACACUGGCCAGGdTdT-3’ [59]. Transfection of a commercially available siRNA with a random nucleotide sequence (scrambled) was used as a control.

The cells were transfected using the RNAiMax (Thermo Fisher Scientific, USA) reagent according to the manufacturer’s recommendations. RNA and protein expression of each of the target gene was evaluated after 72 h of incubation with siRNA.

### 4.13. Molecular Docking

Ligand structures were obtained from the PubChem database (https://pubchem.ncbi.nlm.nih.gov/, access date 01.05.2022) or prepared manually using Avogadro 1.93.0 software and optimized using the OpenBabel 3.0.0 software (http://openbabel.org/, access date 01.05.2022) [37] using the FFE force field with Fastest descent and dE ≤ 5e−6 threshold. Protein structures were obtained from the PDB database (https://www.rcsb.org/, access date 01.05.2024) and optimized using the Chiron service (https://dokhlab.med.psu.edu/chiron/processManager.php, access date 01.05.2024) and Chimera software according to [60]. Molecular docking was performed using the AutoDock Vina 1.1.2 (http://vina.scripps.edu/, access date 01.05.2024). To detect possible alternative binding sites and compare the affinities of the ligands for them, the procedure described in the literature [60] was used. As such, molecular docking was performed in two steps: first, we docked each molecule to the whole receptor as one large binding area to locate potential alternative binding sites, then the coordinates of the docking results were clustered and averaged to give the centers of the binding sites. The grid center coordinates are represented in Table 4. In all cases, and exhaustiveness was set to 8.

**Table 4.**
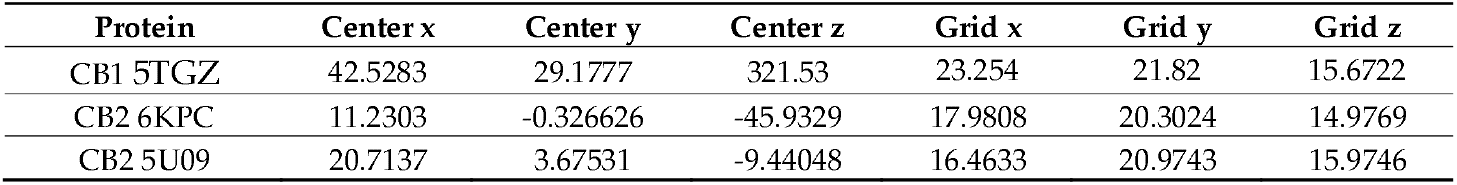
Grid parameters of the docking experiments.

| Protein | Center x | Center y | Center z | Grid x | Grid y | Grid z |
| --- | --- | --- | --- | --- | --- | --- |
| CB1 5TGZ | 42.5283 | 29.1777 | 321.53 | 23.254 | 21.82 | 15.6722 |
| CB2 6KPC | 11.2303 | -0.326626 | -45.9329 | 17.9808 | 20.3024 | 14.9769 |
| CB2 5U09 | 20.7137 | 3.67531 | -9.44048 | 16.4633 | 20.9743 | 15.9746 |

### 4.14. Statistical Analysis

Statistical evaluation was performed using GraphPad Prism 9.3 software. ANOVA with the Holm-Sidak or Dunnett post-test was used to compare the obtained values; p≤0.05 were considered significant.

## 5. Conclusions

In breast cancer cells, the 2-AG analog 2-ADFP induces cell death using an additional two-stage mechanism. At first, it activates CB2 or TRPV1 receptor and induces CREB-dependent COX-2 expression. After that, the accumulated COX-2 oxidizes 2-AG, and its metabolites induce cell death. The observe two-stage mechanism of 2-AG cytotoxicity is an interesting addition to the already known mechanisms of the activity of this compound. On one hand, it could be used as a novel principle for the rational design of anti-cancer compounds. On the other hand, the oxidation products produced by COX-2 may have pro-proliferative activity, and so the discovered COX-2 induction and 2-AG metabolic transformations should be taken into the account during endocannabinoid-based cancer treatment development.

## Author Contributions

Conceptualization, M.G.A. and V.V.B.; methodology, M.G.A. and N.M.G..; investigation, N.M.G., G.D.S., E.I.G., and N.K.; writing—original draft preparation, M.G.A.; writing—review and editing, V.V.B. and N.M.G.; visualization, M.G.A. and G.D.S..; supervision, V.V.B. and M.G.A..; project administration, V.V.B. and M.G.A.; funding acquisition, M.G.A. All authors have read and agreed to the published version of the manuscript.

## Informed Consent Statement

Not applicable.

## Data Availability Statement

The data presented in this study are available on request from the corresponding author. The data are not publicly available due to legal issues.

## Acknowledgments

Not applicable.

## Abbreviations

2-AG: 2-arachidonoyl glycerol
2-ADPG: 2-arachidonoyl 1,3-difluoropropanol
LPI: lysophosphatidylinositol
INT: Iodonitrotetrazolium chloride
LDH: Lactate dehdrogenase
ER: Estrogen receptor
PR: Progesterone receptor
BCIP: 5-Bromo-4-chloro-3-indolyl phosphate
NBT: nitro blue tetrazolium
BCA: bicinchoninic acid

